# Identification of a botulinum neurotoxin-like gene cluster in *Bacillus toyonensis*

**DOI:** 10.1101/2023.07.21.550100

**Authors:** Xin Wei, Briallen Lobb, Kang Wang, Min Dong, Andrew C. Doxey

## Abstract

Clostridial neurotoxins, which include botulinum neurotoxins (BoNTs) and tetanus neurotoxin (TeNT) are the most potent toxins known, and are the causative agents of the neuroparalytic diseases, botulism and tetanus. Until recently, the clostridial neurotoxin family was restricted to the genus *Clostridium*, but members of this protein family have been found in a growing number of non-*Clostridium* species including *Weissella, Enterococcus*, and *Paraclostridium*. Here, we report the bioinformatic identification and analysis of a novel clostridial neurotoxin homolog in a *Bacillus toyonensis* genome recently deposited into the NCBI Genbank database. This putative toxin shares 26-29% identity with its closest BoNT relatives, suggesting that it is likely a novel BoNT-like toxin. It possesses key functional motifs (e.g., HExxH) indicative of toxin protease activity, contains the four characteristic BoNT domains, and is located in a BoNT-like genomic neighborhood containing the upstream non-toxic non-hemagglutinin (NTNH) gene as well as several P47-related genes. Phylogenetically, the toxin clusters as a divergent member of the recently discovered lineage of BoNT-like toxins that includes BoNT/X, BoNT/En, and the insecticidal PMP1. Genomic analysis of the *B. toyonensis* isolate CH177 revealed additional virulence factors and toxin genes indicative of potential pathogenicity targeting an unknown host species. The *Bacillus toyonensis* BoNT-like protein (BTNT) adds to a growing number of non-clostridial BoNT-like toxins, adding further information on the intriguing phylogenetic distribution and evolutionary history of the most potent toxins known.

## Introduction

Botulinum neurotoxins (BoNTs) and tetanus neurotoxin (TeNT) are members of a broader protein family known as clostridial neurotoxins (CNTs), and are the two most potent toxins known [1]. Their sophisticated molecular mechanism of action, role in disease, intriguing phylogenetic distribution and evolutionary history, and use as therapeutics or as scaffolds for design of novel toxin-based therapeutics, make them of considerable interest to the scientific and biomedical community [2–11].

The BoNT mechanism of action can be summarized as follows: after entering host motor neurons by receptor-mediated endocytosis, the BoNT protease is released into the cytosol where it cleaves host SNARE proteins (soluble N-ethylmaleimide-sensitive factor attachment protein receptor). Cleavage of host SNARE proteins inhibits the release of neurotransmitters, resulting in flaccid paralysis [1–3]. The TeNT mechanism of action is similar but it undergoes a unique retrograde transport mechanism to reach inhibitory neurons, resulting in a spastic paralytic phenotype [12].

CNT genes encode a single polypeptide chain that is proteolytically cleaved into two distinct protein chains (light chain and heavy chain), which are connected by an inter-chain disulfide bond. The light chain (LC) includes the N-terminal metalloprotease domain responsible for proteolytic cleavage of host SNARE proteins, and the heavy chain (HC) contains the translocation domain (HN) responsible for translocation of the LC across the host endosome to the cytosol, and the C-terminal receptor binding domains (HCn and HCc) that mediates binding to a variety of host receptors such as polysialogangliosides, synaptotagmin I/II and glycosylated SV2 [13].

Historically, BoNTs were only known to exist within a variety of *Clostridium* species, and were phylogenetically grouped into seven subtypes (A-G). BoNT subtypes, and TeNT which falls within the BoNT phylogenetic tree, are highly divergent from one another, exhibiting amino acid sequence identities as low as ∼30% [14]. Despite their sequence divergence, they are structurally and functionally similar, and retain their overall mechanism of action with the notable exception of TeNT as described above. In addition, different BoNT serotypes exhibit unique host-specificities, with BoNT/A, B, E, and F known to cause human botulism, and subtypes C, D and hybrid CD associated with wildlife botulism outbreaks.

The extremely limited taxonomic distribution, unique structure, and remarkable potency of CNTs, has made them intriguing targets of study for researchers interested in the evolution of toxins and proteins in general [7, 8, 15, 16]. The first indication that BoNT-related proteins might exist outside of the *Clostridium* genus came in 2015 with a study by Mansfield et al. that reported the identification of a BoNT homolog in the organism, *Weissella oryzae* [16]. This highly divergent BoNT-like protein retained several characteristics of BoNTs and was reported to retain the ability to cleave SNARE substrates [16, 17], but lacked several other characteristics of BoNT gene clusters including the presence of nearby accessory genes. The host-specificity of this “BoNT/Wo” protein is still unclear.

Since then, several additional non-clostridial BoNTs have been detected through genome analysis and database searching [18], including BoNT/En from a strain of *Enterococcus faecium* [19, 20] and PMP1 from *Paraclostridium bifermentans* [21]. Phylogenetically, BoNT/En and PMP1 fall into a distinct lineage of BoNT-like toxins, and form their own branch outside of the subfamily of CNTs targeting vertebrates including BoNTs A-G and TeNT. These BoNT-like toxins therefore appear to be a sister clade to the vertebrate BoNTs. Growing evidence suggests that this sister clade of BoNT-like toxins, which also includes the recently discovered BoNT/X from *Clostridium botulinum* strain 111, may have an association with insect hosts [7, 21, 22].

In this work, we report the identification and initial bioinformatic analysis of a novel putative BoNT-like toxin from a particular strain of *Bacillus toyonensis* (“isolate CH177”), whose genome was recently added to the NCBI database on Apr 12, 2023. Our analysis of *B. toyonensis* BoNT-like toxin suggests that it is related to the BoNT/X/En/PMP1 lineage and that it likely forms a novel BoNT-like toxin with unique properties.

## Results and Discussion

### BLAST-based identification of BoNT-related genes in B. toyonensis

On July 6, 2023, we performed a BLASTp search against the NCBI nr database with default parameters using BoNT/A1 (NCBI accession # WP_011948511) as a query. A previously unreported homolog annotated as “hypothetical protein” (accession # HDR7951433) from *Bacillus toyonensis* was detected with an *E*-value of 3 x 10^-53^, and amino acid sequence identity of 26.87%. A second protein (HDR7951432), also from *B. toyonensis*, was detected with an *E*-value of 4 x 10^-33^ and 24.6% amino acid sequence identity. Inspection of both sequences and subsequent BLAST analysis revealed that the first protein (HDR7951433) is a homolog of BoNT, while the second (HDR7951432) is a homolog of the NTNH protein, whose gene is typically encoded upstream of *bont* in *bont* gene clusters. Consistent with this, a zinc-metalloprotease HExxH motif unique to BoNTs was found in HDR7951433 but was absent in HDR7951432. We therefore designated HDR7951433 as BTNT for putative *<u>B</u>acillus toyonensis <u>n</u>eurotoxin* and HDR7951432 as BT-NTNH.

Next, we aligned the putative *B. toyonensis* neurotoxin (BTNT) with other members of the BoNT family (**Figure 1**). It is most similar to PMP1 from *Paraclostridium bifermentans*, but is still highly divergent at only ∼29% identity. This degree of sequence divergence is expected for different serotypes within the BoNT family as shown in **Figure 1**.

**Figure 1.**
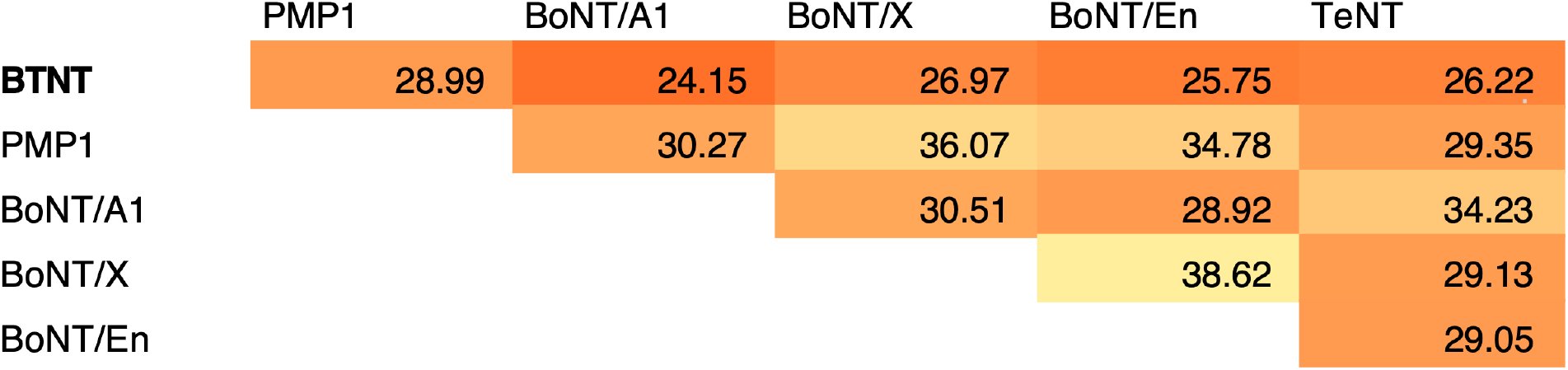
Pairwise sequence similarity of BoNT toxins including the newly identified BoNT-like toxin from *B. toyonensis* (HDR7951433.1).

### Gene neighborhood analysis

To examine the gene neighborhood surrounding the newly identified *B. toyonensis* BoNT-like toxin gene, we retrieved the associated contig (DAOJGQ010000093.1) and gene annotations from the NCBI Assembly (GCA_029709895.1), and visualized it using AnnoView (annoview.uwaterloo.ca). We also manually analyzed and annotated the nearby genes using BLAST. *btnt* has a gene neighborhood structure that is characteristic of other *bont* genes with the *bt-ntnh* gene located immediately upstream as expected, as well as three P47/ORFX-like genes (**Figure 2**). BLAST analysis revealed that the P47/ORFX containing proteins putatively match the proteins ORFX3, P47, and ORFX2 in *P. bifermentans, E. faecium*, and *C. botulinum*. This general gene arrangement is conserved among other members of the X/En/PMP1 clade of BoNT-like toxins, and also some other BoNT genes (e.g., BoNT/A4). It is noticeable that *p47* is located within the *orfX* locus in the genomic neighborhood of BNTN, whereas it is usually located upstream or downstream of the *orfX* locus in the genomic neighborhood of other BoNT toxins. As the available contig is short, the identity of genes further upstream and downstream are currently unknown.

**Figure 2.**
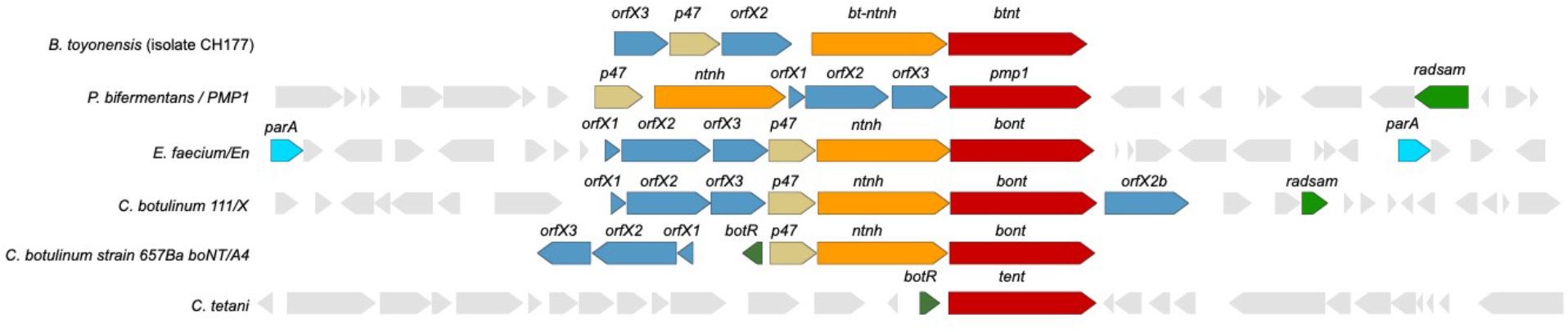
Visualization of a newly identified *bont*-like gene cluster in *B. toyonensis* containing homologs of *bont, ntnh, and* P47-domain containing proteins. Also shown for comparison are *bont* gene clusters from other species.

### Sequence analysis

Next, we aligned BTNT with all representatives of all other BoNT subtypes, and examined the alignment for presence/absence of key functional residues/motifs (**Figure 3**). Although BTNT contains the conserved HExxH motif as described earlier, it appears to lack the double Cys residues within the disulfide linker region at the canonical positions conserved in other BoNTs. This conserved disulfide bond connects the heavy and light chain, and is essential for the toxicity of neurotoxins [23]. However, other Cys residues may perform this role, such as the Cys-590 located eight residues further downstream. BTNT also possesses the conserved PxxG motif identified in the translocation domain of BoNTs and more distantly related BoNT homologs [24]. It lacks the SxWY ganglioside cell receptor-binding motif [25] but possesses a similar SSEY motif at this site, which could indicate an ability to bind a similar receptor.

**Figure 3.**
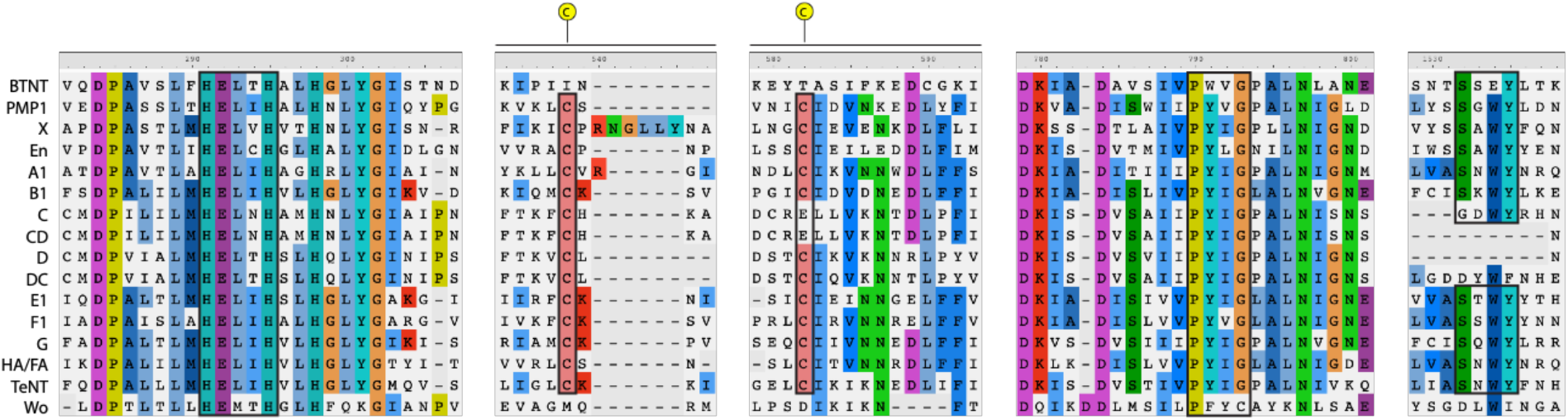
Alignment of *B. toyonensis* BoNT-like toxin with other subtypes and presence/absence of key functional motifs.

### Phylogenetic analysis

To investigate the evolutionary relationships between BTNT and other BoNTs, we constructed a maximum-likelihood tree using IQ-TREE (**Figure 4**). BTNT clustered with the X/En/PMP1 lineage as the outermost branch. Its closer evolutionary relationship to this BoNT lineage is consistent with its gene neighborhood structure (**Figure 2**).

**Figure 4.**
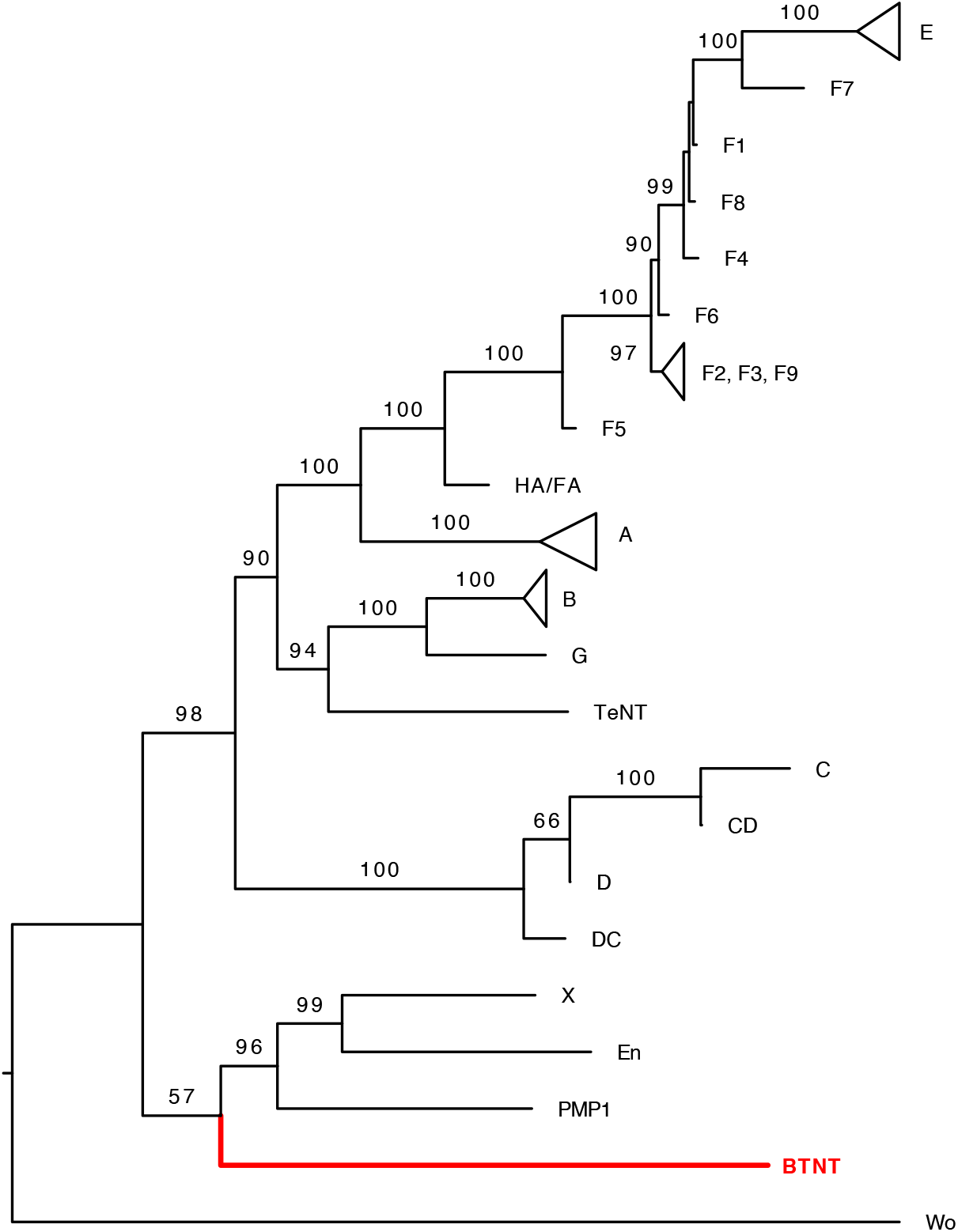
Maximum-likelihood phylogenetic tree of BoNTs and related toxins including the newly identified *B. toyonensis* BoNT-like toxin. Bootstrap values > 50 are indicated above the nodes.

### *Genomic analysis B. toyonensis* isolate CH177

Based on available information in the NCBI, the BTNT-containing *B. toyonensis* strain was recently added to the NCBI database under the accession # GCA_029709895.1 and the isolate name “CH177”. The assembly has 296 contigs, and the draft WGS was generated using the SKESA v2.2. de novo assembler [26].

*B. toyonensis* isolate CH177 was found to be phylogenetically and genomically similar to other *B. toyonensis* strains (data not shown). All currently assembled *B. toyonensis* strains lack detectable homologs of BoNT genes, suggesting that the BTNT gene cluster is a recent acquisition into *B. toyonensis* isolate CH177. Further sequencing and assembly will be required to establish whether it is chromosomally encoded or whether it resides on a plasmid that has been uniquely acquired by this strain.

Next, we examined the CH177 assembly for additional evidence of pathogenicity-related genes using VFanalyzer within the VFDB database [27]. VFanalyzer detected hemolytic enterotoxin genes (*hblA, hblC*, and *hblD*), and a non-hemolytic enterotoxin (*nheC*), all of which have been detected in other related *Bacillus* species. Notably, in a commonly used probiotic *B. toyonensis* strain, these toxin genes are not detectably expressed or are expressed at much lower levels than *B. cereus* [28]. Also detected was the PlcR-PapR quorum sensing system, a hemolysin III family protein, haemolysin *xhlA* gene, and sphingomyelinase. The *islA* gene was also detected, which is a virulence factor in *Bacillus cereus* and has been shown to be expressed in the insect hemocoel and under iron-depleted conditions [29, 30]. Ultimately, these predictions suggest a pathogenicity-related function for CH177, but the host is still unknown.

## Methods

### Sequence and phylogenetic analysis of BoNT-like toxin in B. toyonensis

The BoNT-like toxin in *B. toyonensis*, PMP1 (WP_150887772.1) as well as representative BoNT sequences for each subtype (downloaded from bontbase.org) were used to create a multiple protein sequence alignment. A multiple alignment was then constructed using MAFFT v7.407 with the L-INS-i algorithm [31]. Jalview was used for alignment visualization. IQ-TREE version 2.0.3 was used to generate a phylogenetic tree of all BoNT-toxins [32]. We then midpoint rooted the tree, collapsed some clades that include toxins from the same subtype, and removed bootstrap values lower than 50%.

### Gene neighborhood analysis

The neighboring genes in the ±20kb surrounding the neurotoxin genes were obtained from the NCBI database in .gbk format. These .gbk format files were uploaded to AnnoView (annoview.uwaterloo.ca) for gene neighborhood visualization. Genes in *B. toyonensis* were annotated based on BLAST searches against the NCBI-nr protein database, while all other gene annotations were derived from Wei et al. [22]

### Genome analysis of B. toyonensis strain CH177

A dataset of 310 *B. toyonensis* genomes was downloaded from the NCBI (https://www.ncbi.nlm.nih.gov/datasets/genome/?taxon=155322). A whole genome SNP-based phylogeny was created using Parsnp v1.2 [33] with default parameters and the *B. toyonensis* P18 strain (accession # GCF_016605985.1) selected as a reference genome. The predicted proteome (.fasta file) was submitted to the VFanalyzer tool in the VFDB database [27] with the *Bacillus* genus chosen as a reference taxon. Default parameters were used.

